# Ultra High-Throughput Multiparametric Imaging Flow Cytometry: Towards Diffraction-Limited Sub-Cellular Detection

**DOI:** 10.1101/695361

**Authors:** Gregor Holzner, Bogdan Mateescu, Daniel van Leeuwen, Gea Cereghetti, Reinhard Dechant, Andrew deMello, Stavros Stavrakis

## Abstract

Flow cytometry is widely recognized as the gold-standard technique for the analysis and enumeration of heterogeneous cellular populations and has become an indispensable tool in diagnostics,^1^ rare-cell detection^2^ and single-cell proteomics.^3^ Although contemporary flow cytometers are able to analyse many thousands of cells per second, with classification based on scattering or fluorescence criteria, the vast majority require unacceptably large sample volumes, and do not allow the acquisition of spatial information. Herein, we report a sheathless, microfluidic imaging flow cytometer that incorporates stroboscopic illumination for blur-free fluorescence and brightfield detection at analytical throughputs in excess of 60,000 cells/s and 400,000 cells per second respectively. Our imaging platform is capable of multi-parametric fluorescence quantification and subcellular (co-)localization analysis of cellular structures down to 500 nm with microscopy image quality. We demonstrate the efficacy of our approach by performing challenging high-throughput localization analysis of cytoplasmic RNA granules in yeast and human cells. Results suggest significant utility of the imaging flow cytometer in the screening of rare events at the subcellular level for diagnostic applications.

---

Imaging flow cytometry (IFC) is a hybrid technology which is using advantages of microscopy and cytometry for high-throughput cellular analysis, through rapid imaging of single cells within flowing environments. Such an approach provides for enormous enhancements in information content but is accompanied by a number of technological challenges, such as the ability to acquire high-resolution (blur-free) images of individual cells, integrate multiple imaging modes (e.g. fluorescence, bright-field and dark-field imaging) and achieve adequate detection sensitivities (when using short exposure times).^4^ Commercial imaging flow cytometers (Amnis ImageStream X) have successfully combined the power of microscopy and flow cytometry, to capture images of single cells in flow.^5^ Through the use of hydrodynamic focusing, wide-field sample excitation and time delay and integration charge-coupled device (TDI-CCD) image sensors, the systems offer high resolution and multiplexed imaging capabilities at the expense of low-moderate throughput (3000 cells/sec).^4^ Significantly, such imaging flow cytometers have been used to good effect in the detection and analysis of tumor cells^6,7^ and acute leukaemia^8^. When compared to traditional (single-point) flow cytometry techniques, the imaging capability allows for the detection of chromosomal signalling, antigen localization and other events that occur within the cell.^6,8^ Finally, imaging cells in flow avoids the requirements of membrane staining which is important for performing image segmentation under static conditions.^9^

As noted, a key drawback associated with conventional imaging flow cytometry is the low analytical throughput (between 2000 and 3000 cells per second at 20x magnification), which is more than one order of magnitude lower than non-imaging flow cytometers. To address this limitation and realize high-throughput imaging flow cytometry, recent studies have leveraged the capabilities of microfluidic systems to manipulate, order and process micron-sized objects in a controlled and high-throughput manner. For example, Rane *et al*. recently presented an imaging flow cytometer based on inertial focusing for sheathless manipulation of cells and stroboscopic illumination for fluorescence imaging at high linear velocities, achieving a maximum throughput of 50,000 cells/s.^10^ In addition, Miura and co-workers reported the use of light-sheet excitation of flowing cells within a mirror-embedded microfluidic device. Such an approach was successful in extracting fluorescence images of rapidly moving adenocarcinoma cells at throughputs up to 10,000 cells/s.^11^ Instead of using a (CCD) camera for imaging, images may be reconstructed from photomultiplier tube (PMT) signals through spatial-temporal transformation. For example, *Han et al.* produced high-resolution fluorescence and scatter-based images of rapidly moving cells at a throughput of approximately 1,000 cells/s.^12^ To overcome the problem of preserving image quality when cells are moving at high linear velocities, a free-space pulse-stretching method called free-space angular-chirp-enhanced delay (FACED), has recently been used for time-stretch imaging in the visible-wavelength regime, as well as for fluorescence imaging of rapidly moving beads.^13,14^ Additionally, laser-scanning confocal fluorescence microscopy based on frequency-division multiplexing has been used for high-throughput confocal fluorescence imaging of cells at frame rates of 16,000 frames/s.^14,15^ Recently a deep-learning-assisted image-activated cell sorting platform reported real-time sorting of microalgal and blood cells based on intracellular protein localization.^16^ However, it should be noted that all these approaches, in addition to requiring exotic and complex optical hardware, only report gross morphological features, such as cell/nucleus areas and perimeters. Moreover, the localization of subcellular features equal to or less than 1 micron in size (with microscopy image quality) at throughputs more than 10,000 cells/s has yet to be reported.

Herein, we present a high-throughput imaging flow cytometer capable of multi-parametric imaging (multi-colour fluorescence and bright-field analysis), accurate cell sizing and most importantly subcellular (co-)localization detection of features close to the diffraction limit. To demonstrate utility, we apply the developed platform to the ultra-high-throughput quantitative imaging analysis of cytoplasmic RNA granules in yeast (stress granules) and humans cells (P-bodies). To enable acquisition rates in excess of 10,000 cells/s, we combined stroboscopic illumination, a technique able to generate blur-free images of objects moving at high linear velocities^10^, with elasto-inertial 3D cell focusing^17,18^, to provide for enhanced control of both flow velocities and cell position within the axial depth of focus of the objective lens (Fig. 1a,b, **Supplementary Fig. 1** and **Supplementary Note 1**). The microfluidic device is simple in construction, incorporating a single (90 mm long, 665 µm wide and 59 µm deep) channel containing two switchbacks and an imaging zone located close to the outlet. Multi-parametric detection involves both bright-field imaging, for sizing and morphology measurements, and multi-colour fluorescence detection, for subcellular localization detection (Fig. 1c and **Supplementary Movie 1**). Computational fluid dynamics simulations were used to confirm experimental observations of flow distribution uniformity in the region where the cells enter the microchannel (**Supplementary Fig. 1**). To eliminate motion blur when imaging rapidly moving cells, we stroboscopically manipulated the excitation beam between 10 and 20 µs. This allowed extraction of high-resolution images (**Supplementary Fig. 2**) of cells moving at velocities between 0.03 and 0.05 m/s, with a spatial blur of no more than 500 nm (**Supplementary Fig. 2**). Additionally, light sheet illumination maximizes the excitation efficiency by concentrating photons to a small volume and results in intensities one order of magnitude higher than standard epifluorescence excitation (**Supplementary Fig. 2c**). Using such an approach, we were able to probe the size and fluorescence of labelled Jurkat cells at an experimental throughput of 61,000 cells/sec (**Supplementary Fig. 3**, **Supplementary Note 2**). The ability to resolve different fluorescent intensities and quantitatively measure low-fluorescence sub-populations defines the analytical sensitivity of a flow cytometer. To this end, the system was calibrated with MESF (Molecules of Equivalent Soluble Fluorophores) beads, yielding a detection sensitivity of 3200 MESF units (**Supplementary Fig. 4** and **Supplementary Note 3**). Additionally, adoption of a dual view optics configuration, allowed the implementation of a bright-field imaging modality for the concurrent investigation of cell morphology (**Supplementary Fig. 5**).

**Fig. 1:**
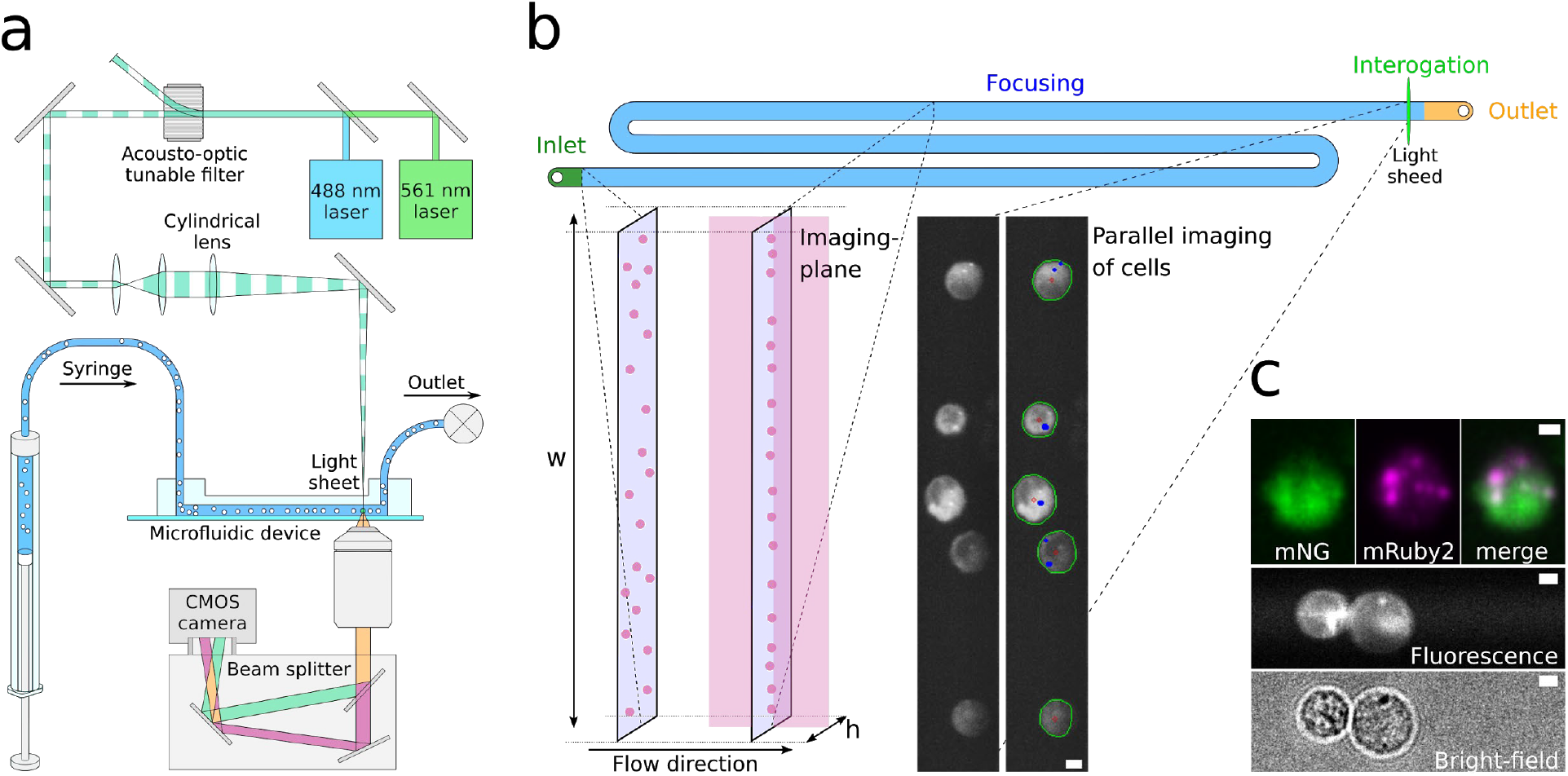
A high throughput multiparametric imaging flow cytometer. (**a**) The imaging flow cytometry platform integrates stroboscopic multi-colour light sheet illumination, a microfluidic cell focusing system, a dual colour beam splitter and a CMOS camera. (**b**) Top view of the microfluidic channel used to position the cells in the imaging plane. The microfluidic device consists of a straight, high aspect ratio channel with one inlet and one outlet. Through the use of a viscoelastic carrier fluid, cells can be precisely focused in the centre plane of the channel. Cells are imaged upstream of the outlet using stroboscopic light sheet illumination, with image processing being used for cell identification and spot foci counting. (**c**) Representative dual colour, fluorescence images of 293T cells labelled with mNG (left, top) and mRuby2 (middle, top) channel. The corresponding merged image is shown in the top, right panel. Images in the lower two panels show the simultaneous acquisition of fluorescence and bright-field images at high-throughput. Scale bars are 5 µm.

To evaluate the performance of the high-throughput imaging flow cytometer in quantifying subtle sub-cellular structures, we analysed the localization of P/GW-bodies and stress granules. P-bodies are small (500 nm - 1 µm) membrane-less cytoplasmic organelles that are intimately involved in RNA metabolism pathways, including RNA decay, stress-induced translational inhibition and RNA silencing.^19,20^ Stress granules are dense cytosolic aggregations that form in response to various stresses (such as starvation or oxidative stress), and are thought to contribute to cellular adaptation by sequestering proteins and RNA to prevent their degradation and allow the efficient restarting of cell growth upon stress relief.^20–22^ First, we chose to study the localization of yeast pyruvate kinase, Cdc19, which aggregates in response to starvation and heat stress (Fig. 2a,b).^23,24^ In line with previous experiments,^24^ analysis of images at 60x magnification indicated that Cdc19-GFP was found uniformly in the cytosol under control conditions, but formed between 1 and 2 granules per cell during the stationary phase (48 hours, long-term glucose starvation). In addition, based on these images, we evaluated the subcellular spatial resolution of the IFC to be ~500 nm (**Supplementary Fig. 6**); a value equivalent to static images ^23^.

**Fig. 2:**
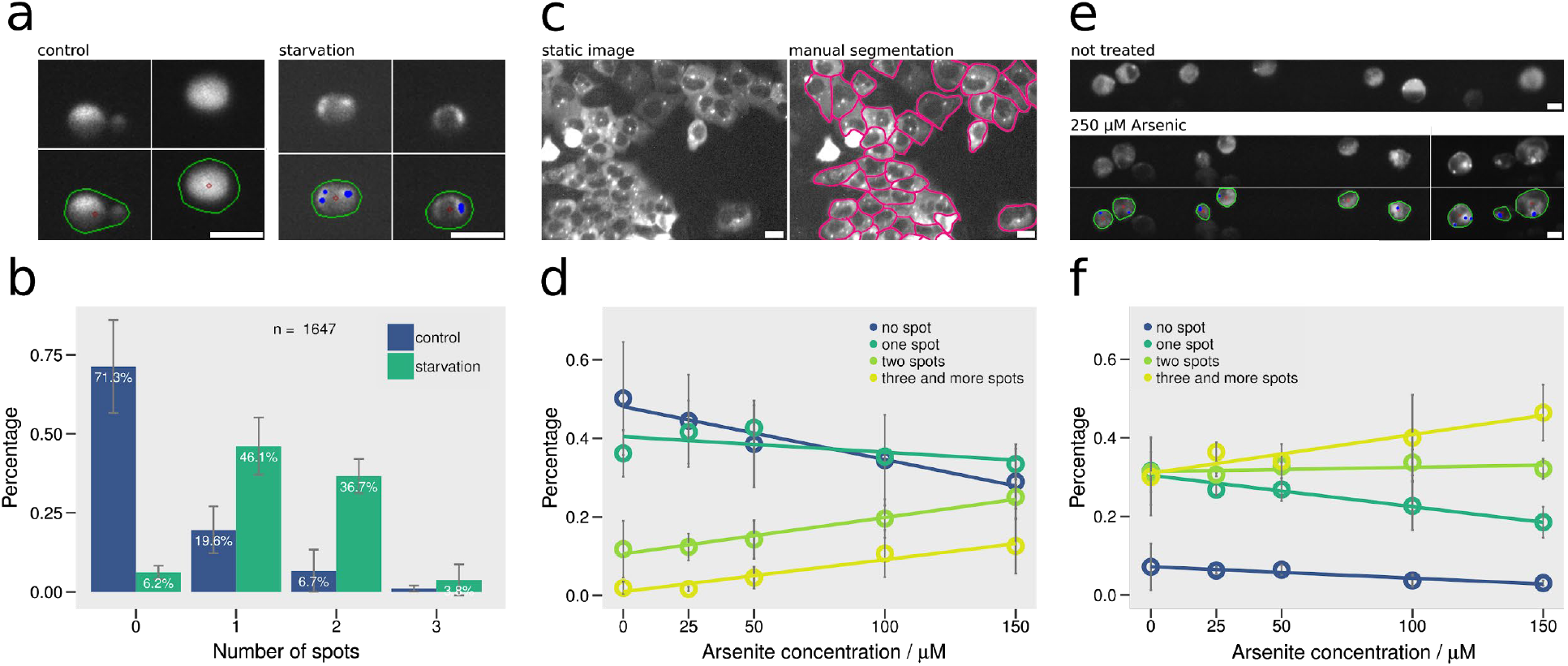
Variation in the number of P-bodies and stress granules in yeast cells in response to external stimuli. (a) Flow cytometry images of yeast growing within the exponential growth phase (control) and within the stationary phase (starvation). Scale bars are 5 µm. (b) Percentage of yeast cells containing given numbers of aggregates, highlighting a clear difference between the control and starvation samples. (c) Adherent 293T cells imaged under static conditions and manually segmented, for mNG-AGO2-positive cytoplasmic granule counting. Scale bars are 20 µm. (d) Percentage of adherent 293T cells containing given numbers of aggregates as a function of arsenite concentration under static imaging conditions: higher arsenic concentrations lead to an increase in the number of mNG-AGO2-positive granules. (e) 293T cells images obtained in the imaging flow cytometer at 15x magnification are automatically segmented and processed. Scale bars are 10 µm. (f) In the case of flowing cells, the mNG-AGO2-positive granule number increases with the increase of the arsenic concentration showing the same trend with the static image in (d).

We next investigated the sub-cellular localization of Argonaute 2 (AGO2) and Trinucleotide Repeat-Containing Gene 6A protein (TNRC6A), two directly-interacting proteins from the RNA silencing pathway and P-body constituents in human cells.^25,26^ As previously reported^27^, 293T cells stably expressing mNeonGreen-tagged AGO2 (mNG-AGO2) exhibit both a diffuse cytoplasmic emission pattern along with 1-3 granular cytoplasmic structures per cell (characteristic of P/GW-bodies) and that are absent in control cells expressing mNG only (Fig. 2e). Several stress-activated kinase pathways known to be activated in cancers (e.g. p38, AKT) directly control Argonaute shuttling and transfer into P/GW-bodies (as well as stress granules) by direct phosphorylation.^28,29^ Among stresses, oxidative stress induced by arsenite treatment has been previously shown to increase the number of AGO2-positive RNA granules in cells, by triggering the formation of AGO2-positive stress granules.^30^ Accordingly, treatment of cells with arsenite, leads to a dose dependant increase in the number of mNG-AGO2 positive granules per cell, as detected by static fluorescence microscopy (Fig. 2d) and by high throughput IFC with similar trends (Fig. 2e,f). Of note, the substantially distinct cell morphologies for both imaging approaches (i.e. flat versus spherical cells, manual versus automatic segmentation) could explain the difference in absolute counts of AGO2-positive granules. Critically, our approach affords reliable detection and automated quantification of sub-micron subcellular variations (~1 µm, **Supplementary Fig. 6**) in single mammalian cells at throughputs of 20,000 cells/s and at 15x magnification.

Finally, we assessed the possibility of performing dual-colour co-localization assays using high throughput imaging flow cytometry in mammalian cells. We investigated the subcellular localization of both TNRC6A and AGO2 in 293T cells during oxidative stress response. For that purpose, we co-expressed mRuby2-TNRC6A or mRuby-AGO2 in 293T cells stably expressing mNG-AGO2 or mNG (Control). Accordingly, mRuby-TNRC6A (Fig. 3a, **panels 7-8**), or mRuby-AGO2 (Fig. 3a, **panels 3-4**), also form cytoplasmic foci co-localizing with mNG-AGO2 (but not with the control mNG), as demonstrated by quantification of granule co-localization percentage (Fig. 3b) and pixel intensity correlation (Fig. 3c, **Supplementary Fig. 7**). Importantly, these observations confirm our ability to detect subtle subcellular localization events (of distinct proteins) within the same cell at high resolution and at throughputs of 20000 cells/s (15x magnification), 10x faster than the commercial ImageStream imaging flow cytometer

**Fig. 3:**
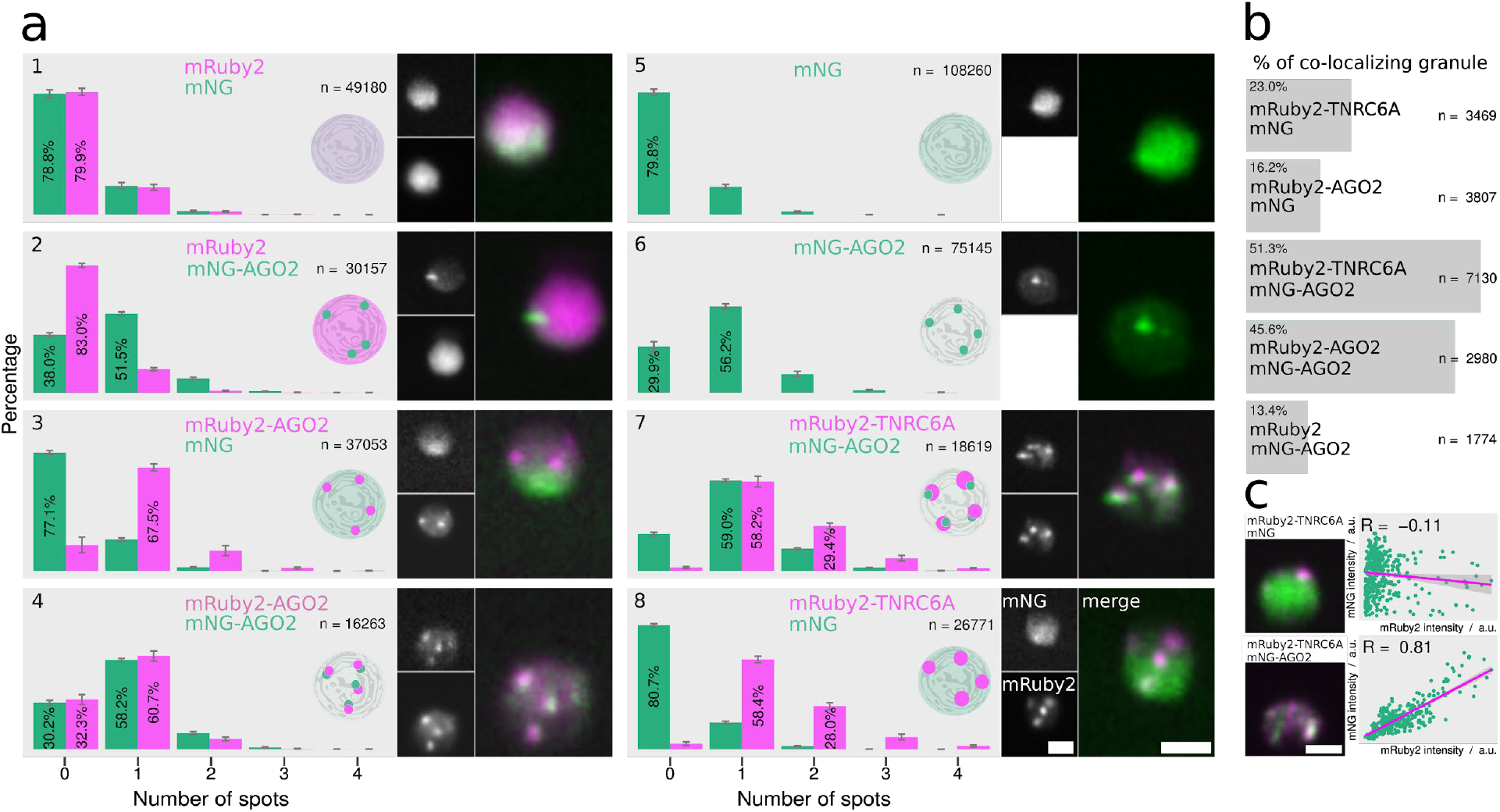
Dual color IFC for high-throughput co-localization studies. 293T cells expressing mNG and mNG-AGO2 are transfected with different mRuby-tagged constructs. (**a**) Granule counts for green and red channels, with the y-axis representing percentages of fluorescently labelled cells. Cartoons illustrate representative localization states of the corresponding transfected cells. Small image panels show green (top) and red (bottom) channels, with an overlay of both channels being provided in the large image on the right-hand side. Scale bars are 10 µm. (**b**) Percentage of granules occupying the same spot in both channels. (**c**) Overlay images (left panels) and scatter plots (right panels) of mNG and mRuby2 pixel intensities for cells co-expressing mRuby2-TNRC6A/mNG (top) and mRuby-TNRC6A/mNG-AGO2 (bottom). Pearson’s correlation coefficients and associated p-values indicate a weak correlation for the mRuby2-TNRC6A/mNG cell, while the mRuby-TNRC6A/mNG-AGO2 cell shows a strong correlation for both the red and green channel.

To conclude, we have presented a new microfluidic platform for high-resolution, image-based analysis of cells at ultra-high-throughputs. Such a single-cell analysis capability leverages and combines the high-throughput nature of conventional flow cytometry with the enhanced sensitivity and information content associated with optical microscopy. Critically, the adoption of a wide-channel microfluidic format, facilitates system automation and greatly simplifies the optical detection scheme. As we have shown, such advances allow the realization of high-throughput cell sorting and provide for an increase in analytical throughput of several orders of magnitude when compared to existing imaging flow cytometers. Indeed, as shown in **Supplementary Fig. 8** and **Supplementary Note 4**, the platform is successful at processing mixed populations of Jurkat and Human B-lymphoid cells at rates in excess of 400,000 cells/second.

## METHODS

### Microfluidic Device Fabrication

Microfluidic devices were fabricated using standard soft-lithographic techniques. Briefly, the two-dimensional channel pattern was designed using AutoCAD (AutoCAD 2017, Autodesk,CA,USA) and printed onto a transparent film photomask (Micro Lithography Services Ltd, Chelmsford, United Kingdom). This photomask was subsequently used to pattern an SU-8-coated silicon wafer (Microchem Corporation, Westborough, USA) using conventional photolithography. A 10:1 mixture of polydimethoxysilane (PDMS) monomer and curing agent (Sylgard 184, Dow Corning, Midland, USA) was poured over the master-mold and peeled off after polymerization at 70°C for 4 hours. Inlet and outlet ports were created using a hole-puncher (Technical Innovations, West Palm Beach, USA) and the structured PDMS substrate then bonded to a planar glass substrate (Menzel-Glaser, Braunschweig, Germany) after exposing both surfaces to an oxygen plasma (EMITECH K1000X, Quorum Technologies, East Sussex, United Kingdom) for 60 seconds.

### Microfluidic Device Structure

A schematic of the microfluidic device, consisting of a straight high aspect ratio channel, is shown in Fig. 1b. Channel cross-sections were 59 × 665 µm (for experiments using a 20X objective), 59 × 1331 µm (for experiments using a 10X objective) and 38 × 332 µm for experiments using yeast cells. These dimensions guaranteed an even distribution of cells across the channel, as shown in **Supplementary Fig. 1** and **Supplementary Fig. 6.** The height of the microfluidic channel was approximately 59 microns in order to satisfy the blockage ratio criterion, *β*=*a*/*h*, (where a and *h* are the cell diameter and the characteristic height of the microchannel) to be smaller than 0.25. The average diameter of the mammalian cells used in this study was 13 µm and the average diameter of yeast cells is approximately 4 µm. (Such conditions yielded single file/plane focusing.^17^ Imaging region correspond to different areas of the CMOS sensor (1335 µm × 1335 µm) and are defined by the magnification of the objective lens (**Supplementary Table 2**).

### Device Operation

PDMS devices were rinsed in water and then incubated for 10 minutes in a 2% w/w Pluronic ® F-127 (Sigma-Aldrich, Buchs, Switzerland) solution to prevent cells sticking to the channel walls. The cell suspension (with concentrations up to 25 million cells per mL) was loaded into a 1 mL syringe (Gastight Syringes, Hamilton Laboratory Products, NV, USA) and delivered at the desired flow rate using a precision syringe pump (neMESYS, Cetoni, Korbussen, Germany). The microfluidic device was placed on a motorized *xy* translation stage (Mad City Laboratories, Maddison, USA) that is mounted on an inverted Ti-E microscope (Nikon, Zurich, Switzerland). In all experiments, cells and beads were suspended in a viscoelastic polyethylene oxide solution (PEO) solution to allow for elasto inertial focusing. Multiple parameters must be controlled to ensure efficient focusing cells into a single file. These include the molecular weight of the polymer, concentration of the polymer solution, microfluidic channel geometry, the blockage ratio and flow rate of the suspension. A detailed description of the specific influence of each of these parameters can be found elsewhere^17^. Based on this, current experiments were carried out in a 500 ppm/1 MDa PEO and 1000 ppm/1 MDa PEO (Sigma-Aldrich, Buchs, Switzerland) solutions for cells and yeast/beads, respectively. A stock solution at a concentration of 10,000 ppm was prepared and aged at room temperature for a month, to ensure stability.^17^ The stock solution was diluted with DPBS (Life Technologies, Zug, Switzerland) to the desired concentration and used immediately or stored at 4 °C.

The cell suspension was loaded into a 1 mL syringe (Gastight Syringes, Hamilton Laboratory Products, NV, USA) and delivered at flow rates of up to 240 µL/minute using a precision syringe pump (neMESYS, Cetoni, Korbussen, Germany). Settling of cells within syringes was minimized by matching the density of the medium to the cell suspension using a 36 % v/v Optiprep Density Gradient Medium (Sigma-Aldrich, Buchs, Switzerland). A commercial calibration kit (Quantum Alexa Fluor 488 MESF, Bangs Laboratories, Indiana, USA) consisting of five microsphere populations surface labeled with increasing amounts of a specified fluorochrome, was used for intensity calibration measurements (**Supplementary Fig. 4**). The volumetric flow rate of the cell suspension was adjusted, so that cells are over-sampled during the imaging process and to receive blur-free images according to relations presented in **Supplementary Fig. 1**. Since the cropping area is slightly larger than the diameter of the cell, oversampling help us to acquire cells in more than 1 frame and thus the identification of its size and content can be precisely defined.

This is achieved by synchronizing the cell velocity to the camera acquisition rate. The image acquisition rate was set to between 2000 and 3000 frames per second, depending on the magnification of the objective and the corresponding size of the region of interest. Accordingly, each cell traverses the microfluidic channel at a time lower than frame rate, and thus will be imaged more than once during the acquisition process. This ensures, no cell passing through the detection region will not be imaged. Using such criteria, and for the current system, cells should move at a linear flow velocity of between 0.03-0.05 m/s, depending on magnification. Crucially, the number of cells detected in each frame is maximized by working at cell concentrations high enough to ensure compact packing.^17^

### Cell culture, transfection and fixation

Experiments were performed on four different cell lines: JURKAT Clone E6-1 (LGC Standards GmbH, Wesel, Germany USA), Human B-lymphoid (Sigma-Aldrich, Buchs Switzerland), 293T Flp-in T-REX (Life Technologies, Zug, Switzerland) and yeast. Jurkat and Human B-lymphoid cell lines were initially tested for mycoplasma contamination and then cultured in RPMI-1640 medium (Life Technologies, Zug, Switzerland) supplemented with 10% (v/v) FBS (Life Technologies, Zug Switzerland) and 1% (v/v) Penicillin-Streptomycin (10,000 U/mL, Life Technologies, Zug Switzerland) in a CO_2_ incubator (New Brunswick™ Galaxy® 170 S, Eppendorf, Schönenbuch Switzerland) at +37°C, 5% CO_2_, according to standard protocols. All experiments were performed on cells in the exponential (log) phase of growth. Cells were fixed in 4% paraformaldehyde for 10 minutes at room temperature, rinsed 3 times in HBSS buffer and incubated with Alexa Fluor 488 WGA (1:200, A12379, Life Technologies, Zug, Switzerland) for 10 minutes at room temperature. After rinsing twice in HBSS, cells were transferred in PBS and mixed with OptiPrep ® and the PEO solution to the desired ratio and used directly.

293T Flp-in T-REX cells expressing mNG or mNG-AGO2 cells were cultured in DMEM medium (Life Technologies, Zug, Switzerland), supplemented with Glutamax and HEPES (Life Technologies, Zug, Switzerland), and 10% fetal bovine serum (FBS, Life Technologies, Zug, Switzerland). It should be noted that our mNG tags also have a 3xFlag and SBP affinity peptide sequence at the N-terminal. Two days before analysis, 8 million cells were seeded in a 10 cm plate in the presence of doxycycline (2 µg/ml) to induce expression of the tagged protein. Transfection of the plasmid was performed using PEImax (Polysciences Europe GmbH, Hirschberg an der Bergstrasse, Germany), by mixing 180 µl of PEImax (1 mg/ml) with 9 µg of plasmid 24 hours after seeding, and replacing the medium after 8 hours. Then, 48 hours after plating, cells were treated with sodium arsenite (Sigma-Aldrich, Buchs Switzerland) for 2 hours. The medium was subsequently replaced with PBS before 8-10 representative images (green, red and phase contrast) for each condition were acquired on a Floid fluorescence microscope (Life Technologies, Zug, Switzerland). Afterwards, adherent cells were trypsinized, and the corresponding cell suspension washed once in PBS, before fixation in 2% paraformaldehyde for 10 minutes. Finally, the cell suspension was washed twice with PBS before re-suspending in DMEM supplemented with 10% FBS, and then washed with PBS.

Yeast media and growth conditions of cells were as described previously.^24^ Briefly, saturated overnight cultures were diluted into fresh synthetic complete media containing 2% glucose for 5 hours (mid-exponential phase), or for an additional 48 hours to ensure entry into the stationary phase (starvation). Cells were fixed using a 3.7% formaldehyde solution and incubated at room temperature for 10 minutes. Cells were washed twice in PBS by centrifugation and then imaged. The yeast strain used in the current experiments was ySS12^24^ (genotype: BY4741, CDC19::CDC19-GFP-HIS3, Life Technologies, Zug, Switzerland)

### Optical Setup & Data Acquisition

The optical system (Fig. 1a) consists of an inverted microscope (Eclipse Ti-E, Nikon, Zurich, Switzerland) equipped with a motorized stage (Mad City Labs, Maddison, USA) and a Dual-View detection system (Cairn Research, Kent, UK). The outputs of a green (561 nm, Coherent Genesis MX, Glasgow, UK) and blue (488 nm, Coherent Genesis MX, Glasgow, UK) laser were combined using a set of mirrors. After transmission through an acousto optical tunable filter (AOTF nC-400-650-TN, AA Opto-electronic, Orsay, France) connected to a RF driver (AA Opto-electronic, Orsay, France), the combined beam was focused to a line with a width approximately equal to the average cell diameter (~15 µm) using a cylindrical lens (LJ1558RM-A, Thorlabs, Lübeck Germany). Yeast experiments required the use of an acylindrical lens (AYL2520-A, Thorlabs, Lübeck Germany) to generate a line width of 5 µm.

A variety of objective lenses, including a 10x (Plan Fluor 10x, NA 0.5, Nikon, Zurich, Switzerland), a 20x (Plan Apo 20x, NA 0.50, Nikon, Zurich, Switzerland) and a 40x (Plan Fluor 40x, NA 0.75, Nikon, Zurich, Switzerland) were used. Additionally, an extra lens that provided 1.5x magnification at the output port of the microscope was also used on occasion. Fluorescence originating from individual cells was collected by the objective lens and passed through the Dual-View optical configuration to obtain dual colour images. Simultaneous brightfield and fluorescence imaging was accomplished using a red LED light source (Spectra X, Lumencor, Beaverton, USA) in combination with the blue laser at 488 nm, as depicted in **Supplementary Fig. 4**.

The Dual-View optical configuration was mounted between the tube lens of the microscope and a CMOS camera (ORCA-flash 4.0, Hamamatsu, Solothurn, Switzerland or Prime 95B™ Scientific CMOS Camera, Roper Scientific, Planegg, Germany) and comprised two mirrors, a dichroic mirror and two emission filters. For dual colour experiments, a dichroic mirror (560 DCXR, AHF, Tubingen, Germany) was used to split the emission light into two colours. Two emission filters D520/35 (AHF, Tubingen, Germany) and LP591 (AHF, Tubingen, Germany) were used to detect mNeonGreen and mRuby2 fluorescence, respectively. The two fluorophores were imaged simultaneously in different regions of the CMOS camera and commercial software (MicroManager 1.4.15, University of California, San Francisco, USA) used to record the images. For brightfield experiments, the longer wavelength channel was used without an emission filter. Synchronization of both the laser and the LED light source with the camera was accomplished using two ESIO AOTF controllers (ESImaging, Kent, UK). The AOTF and the LED were triggered by the CMOS camera with a TTL pulse (matched to the exposure time of the camera) used to achieve synchronization between the laser (SPECTRA X light engine, Lumencor, USA) and the camera. The sampling frame rate of the ORCA-flash 4.0 CMOS camera was 2000 fps, and the AOTF driver generated a pulse length that varied between 13 and 20 µs (depending on the flow rates used) for both laser lines. The laser beam was modified into a slit-shaped profile, oriented orthogonally to the flow direction, through the use of a cylindrical lens. Using this excitation profile, only a narrow field needs to be illuminated. This contrasts with traditional epi-illumination excitation formats, in which the entire field of view is illuminated. The light density provided by the cylindrical lens increases the effective illumination intensity by one order of magnitude when compared to normal epi-illumination modalities.

### Image Processing and Data Analysis

Post-processing of data is performed using algorithms that analyse fluorescence and bright-field images to extract the size, location and the intensity of cells and intracellular features. The image processing algorithm was implemented in Python 2.7 and OpenCV 2.4. The images acquired (and stored as a TIFF stack) were analysed for various cellular attributes, including cell area, cell size, cell position, spot number, spot size, spot location, and fluorescence intensity. Data obtained from the processing algorithm were then plotted as scatter plots and histograms. Initially, the image is read from the TIFF stack and a “de-noising” algorithm implemented in Python to perform intensity-based filtering of noise from the image.^31^ This step is important as it reduces background noise and makes subsequent binary thresholding more reliable. An inbuilt OpenCV function ‘find*Contours*’ is then used to identify cell and spot borders, with each of the cell and spot images being stored as a closed contour. This closed contour provides cell and spot morphology parameters such as cell area, cell size and centroid coordinates. Using the cell border information, the raw image is analysed for fluorescence. The built in function ‘*pointPolygonTest*’ allows assignment of spots to cell contours. The code searches for the maximum fluorescence intensity inside the defined cell area and also computes the mean fluorescence intensity of the cell.

The analysis of all images in a stack, gives a data matrix containing information on the size, area, cell centroid and fluorescence intensity of the cell. These raw data are then filtered to remove artefacts due to image noise and cell debris. The filtered data are then presented in the form of scatter plots and histograms. The CVs (coefficients of variation) were defined as the ratio of the standard deviation to the mean for a measured population.

For multi-parametric detection, it is essential to correlate co-localized events (for example, mRuby2-Ago2 and mNG-Ago2). This is achieved using the information derived from the coordinates of the centroid of the detected event. First data obtained from different detection channels are corrected for offsets in the position of detected event in the image. Subsequently, images from both channels are analysed for cell or spot attributes. Detected events are then correlated by comparing the coordinates of the detected events in both channels. Once the events are correlated, plots of the chosen attributes can be generated.

### Flow cytometry and analysis

A commercial flow cytometer (Astrios MoFlo, San Jose, USA). was used for measuring the coefficient of variation (CV) of fluorescence intensity of Jurkat cells. Data were analysed using R (www.r-project.org). A plot of forward scatter against side scatter was first used to gate cellular events and eliminate events caused by cell debris. A plot of peak area against peak height for the given detection channel was used to gate single cells. Finally, fluorescence intensity histograms were created for the cell population. The number of gated events corresponds to 20,000 cells for statistical robustness in data analysis.

## ACKNOWLEDGMENTS

The authors would like to acknowledge support from the Swiss National Science Foundation and ETH Zürich.

## AUTHOR CONTRIBUTIONS

S.S, G.H and A.dM conceived the project and devised the research plan. S.S and G.H developed the instrumental platform, methodology and performed all experiments, G.H performed image processing, with B.M, D.V.L, G.C and R.D developing the cell protocols and culturing the biological samples, S.S, C.H, B.M and A.dM wrote the manuscript.

## COMPETING FINANCIAL INTERESTS

The authors declare no competing financial interests.

## DATA AVAILABILITY

The data that support the plots within this paper and other findings of this study are available from the corresponding author upon reasonable request.

