## Supplementary Information for "Ultra High-Throughput Multiparametric Imaging Flow Cytometry: Towards Diffraction-Limited Sub-Cellular Detection"

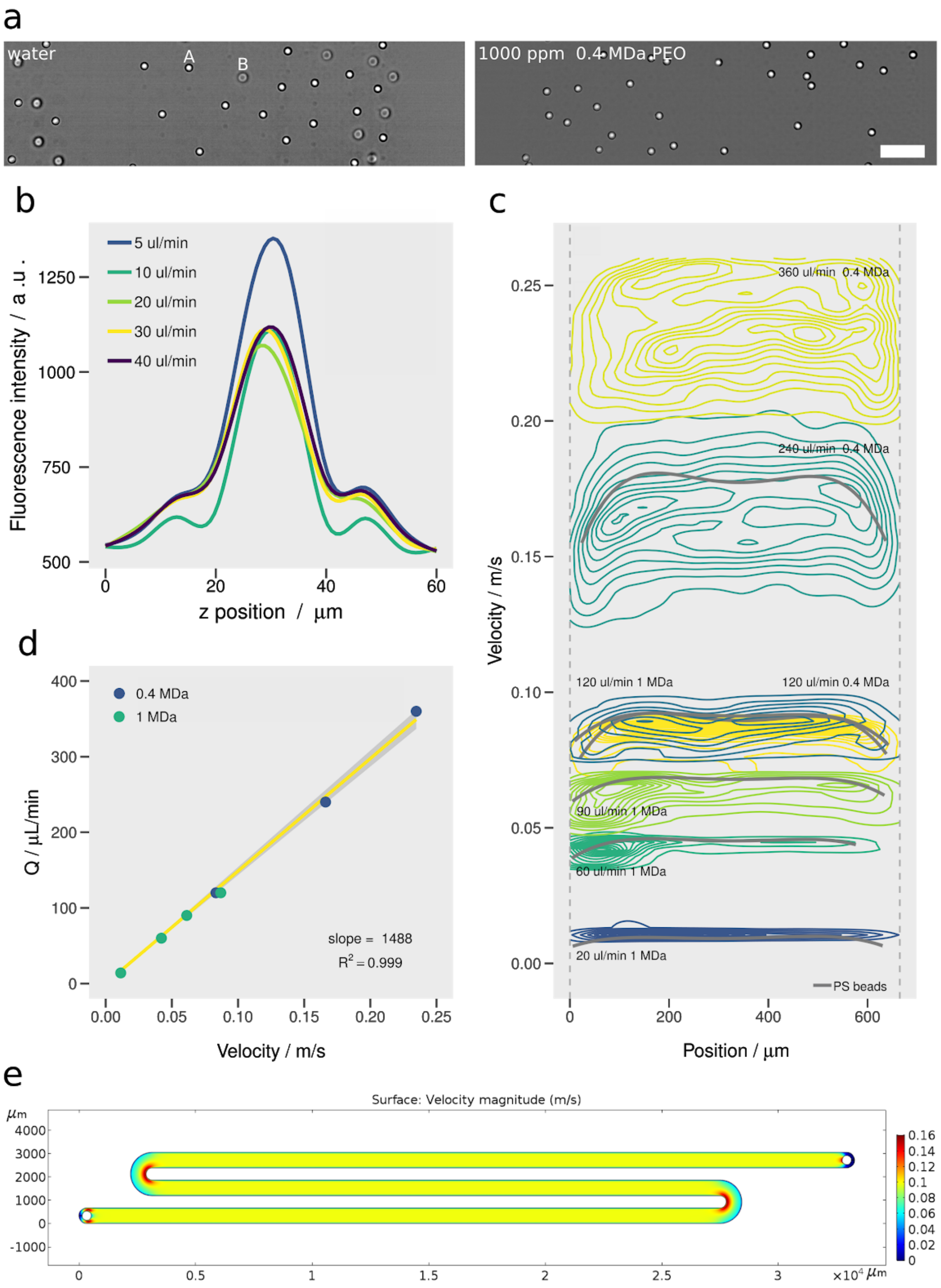


**Supplementary Fig. 1: Elasto-inertial cell focusing**. (a) Brightfield images of polystyrene beads dispersed in water and a 1000 ppm PEO solution. Left panel: 10 µm PS beads dispersed in DI water and flowing at a flow rates of 360 µL/min are focused into two planes (labelled A and B). This behaviour is typical for inertial focusing regimes.[1] Right panel: 10 µm PS beads dispersed in 1000 ppm 0.4 MDa PEO flowing at a flow rate of 360 µl/min are focused to the centre plane of the microchannel. (b) Variation in fluorescence intensity as a function of z position for Jurkat cells stained with Alexa Fluor 488 WGA within a 59 x 665 µm cross-section microchannel. Cells are dispersed in a 500 ppm PEO (1 MDa) solution, with intensities, measured using a spinning disk confocal microscope (Yokogawa W1, Tokyo, Japan). Inspection of the intensity profile confirms that cells are positioned at the centre of the channel cross section. (c) Flow profiles of cells suspended in either 0.4 MDa, 500 ppm PEO and 1 MDa, 500 ppm PEO and moving at velocities between 0.01 and 0.25 m/s through a 55 x 665 μm cross-section microchannel. For all flow rates, the corresponding velocity profiles are homogeneous across the channel, and as expected slightly bended towards the walls of the channel. Flow profiles were obtained from brightfield measurements. (d) Variation of velocity as a function of volumetric flow rate within a 55 x 665 um cross-section microchannel. (e) A two-dimensional COMSOL simulation of the flow velocity throughout the microfluidic device. The observed uniformity of flow profile favours an even cell distribution along the channel and is confirmed by the data shown in (c). The CV of the mean velocity along the detection channel is negligible (<0.99 %), confirming a uniform flow distribution across the entire microfluidic device.

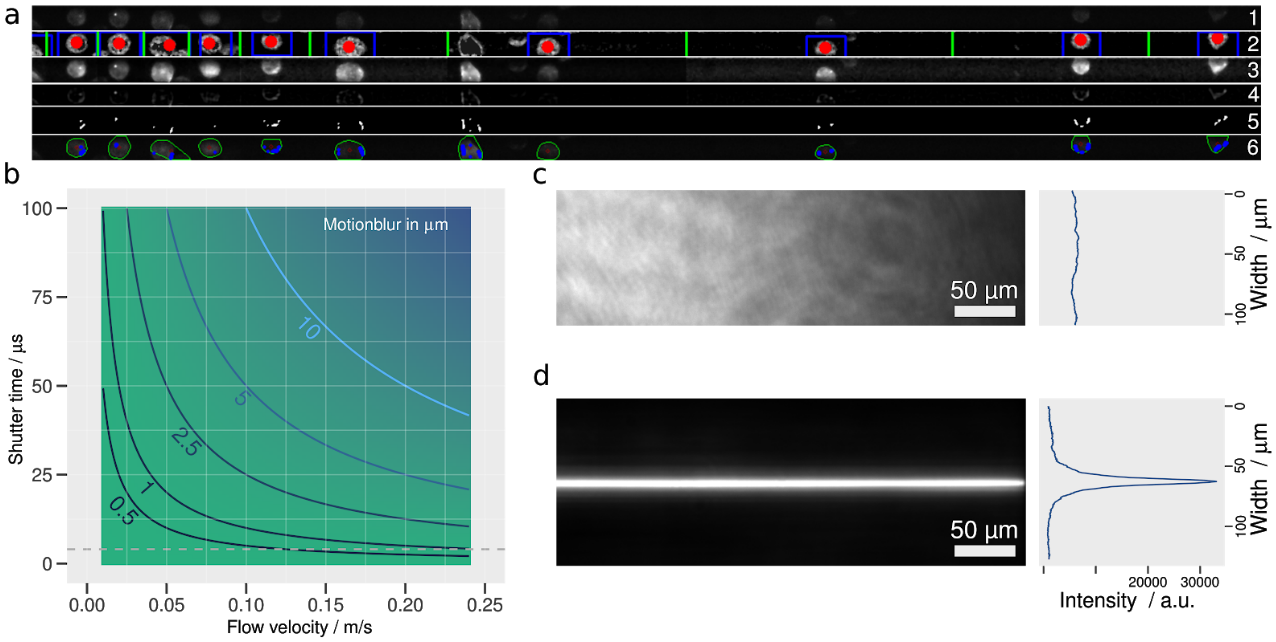


**Supplementary Fig. 2: Evaluation of the motion blur and characterization of the light-sheet illumination.** (a) Description of the image-processing algorithm used to extract contours and related properties of imaged cells. The numbers on the right-hand side indicate the component steps of the image processing workflow: (1) Raw image, (2) Division of frame into multiple regions of interest for contrast adjustment, (3) Contrast adjusted image, (4) Image thresholding to extract cell contours, (5) Identification of intracellular structures and (6) Final image featuring cell and structure contours. (b) Relationship between the illumination pulse duration, flow velocity and motion blur. (c) Cross sectional profiles of epifluorescence (top) and light-sheet illumination (d) using a 100 µM fluorescein solution. An increase in fluorescence intensity of over one order of magnitude is observed for the case of light-sheet excitation (FWHM of ~5 µm) compared to epifluorescence excitation.

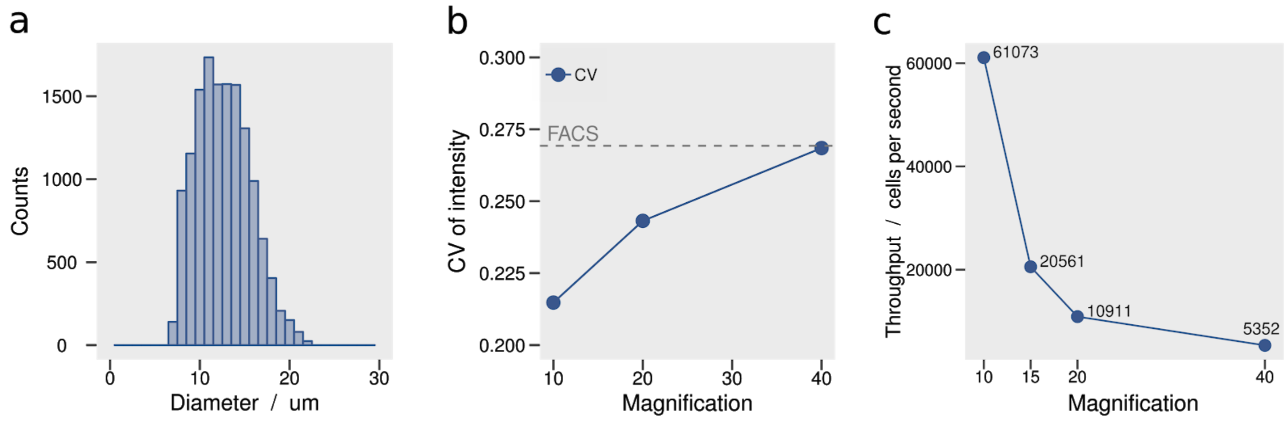


**Supplementary Fig. 3: Performance of the imaging flow cytometer.** (a) Size distribution of Jurkat cells stained with Alexa Fluor 488 WGA, imaged with a 10X 0.5 NA objective. (b) Coefficient of variation for measured fluorescence intensities at three different magnifications: 10X 0.5 NA, 20X 0.5 NA and 40X 0.75 NA. It should be noted that the CV for the 40x 0.75 NA objective is similar to that obtained when using a commercial FACS system (Astrios MoFlo Beckman Coulter, Brea, USA), demonstrating that fluorescence variations are reduced when using a higher magnification objective. (c) Variation in analytical throughput as a function of magnification. An experimental maximum throughput of 61000 cells/second was achieved when using a 10X 0.5 NA objective at a frame rate of 4888 fps.

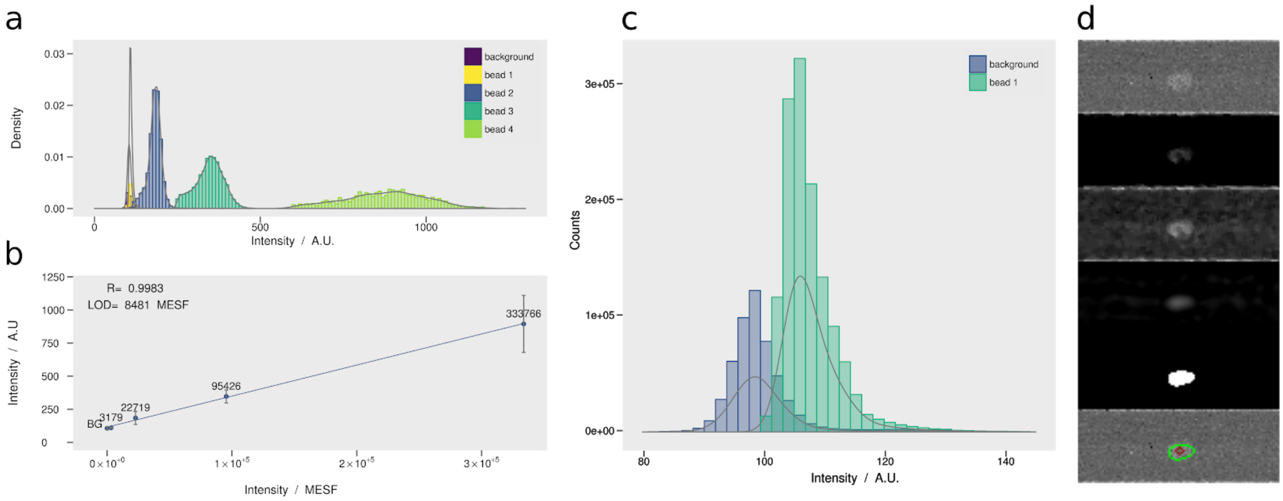


**Supplementary Fig. 4: Evaluation of the analytical sensitivity of the imaging flow cytometer.** (a) Fluorescence Intensity histogram for four microsphere populations surface labeled with increasing amounts of a specified fluorochrome (Quantum Alexa Fluor 488 MESF, Bangs Laboratories, Indiana, USA ). The sample contains four particle sub-populations corresponding to four fluorescence intensity levels. The vendor-specified bead intensities are as follows: bead 1 = 3,179, bead 2 = 22,718, bead 3 = 95,426, and bead 4 = 333,766 MESF units. (b) Fluorescence calibration reporting the measured average intensity as a function of the specified MESF units. The limit of detection per single pixel is calculated to be 8481 MESF units, following the method described by Armbruster and Pry.[2] (c) The lowest intensity beads (bead 1 = 3,179 MESF) can be discriminated after several image processing steps, as shown in (d), despite the fact that calculated LOD is higher than the average bead intensity.

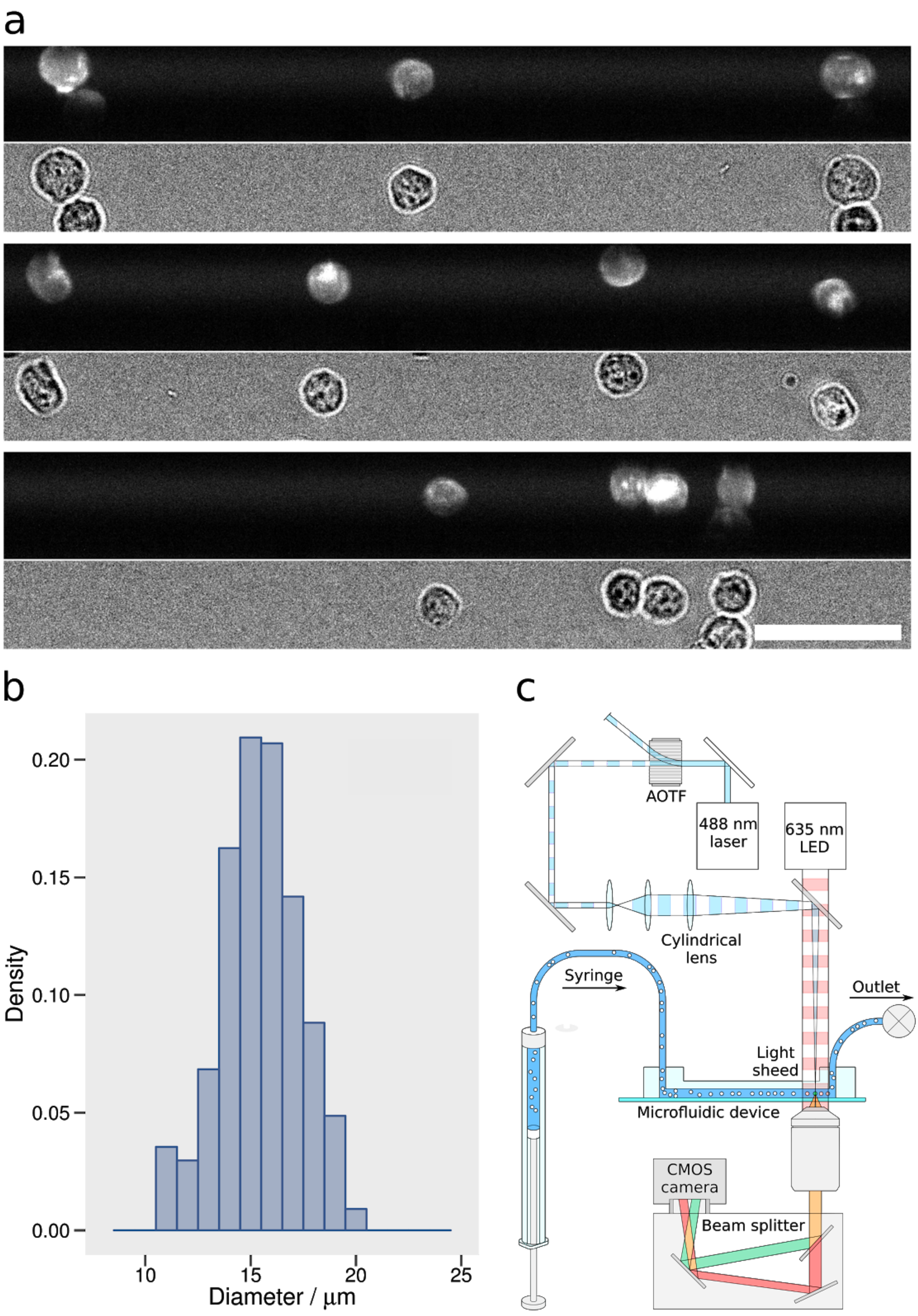


**Supplementary Fig. 5: Multi-parametric detection: brightfield and fluorescence imaging.** (a) Jurkat cells labelled with Alexa Fluor 488 WGA were imaged with a 20x 0.5 NA objective in a dual view configuration, resulting in fluorescence images in the top half and simultaneous bright-field images in the bottom half. The high-resolution fluorescence and bright-field images confirm that cells are tightly focused to the centre plane of the microchannel. Scale bar is 50 μm. (b) Cell size distribution extracted from the brightfield images. (c) Schematic of the optical setup used for fluorescence/brightfield dual imaging. Excitation beams of 488 nm (Sapphire 488 LP, Coherent). The 488 nm beam was passed through an acousto-optic tunable filter, expanded and collimated using a telescope and focused through a cylindrical lens into the microfluidic channel after being reflected by a polychromatic dichroic mirror DM (630DRLP AHF). Collected photons were spectrally filtered using filter (488LP, AHF) and imaged onto a fast CMOS camera. A red LED light source at 635 nm (Spectra X, Lumencor, Beaverton, USA) was used for brightfield stroboscopic illumination. Synchronization of the strobe (excitation) pulse with the integration period of the camera is achieved by using the output signal of the camera to trigger the AOTF.

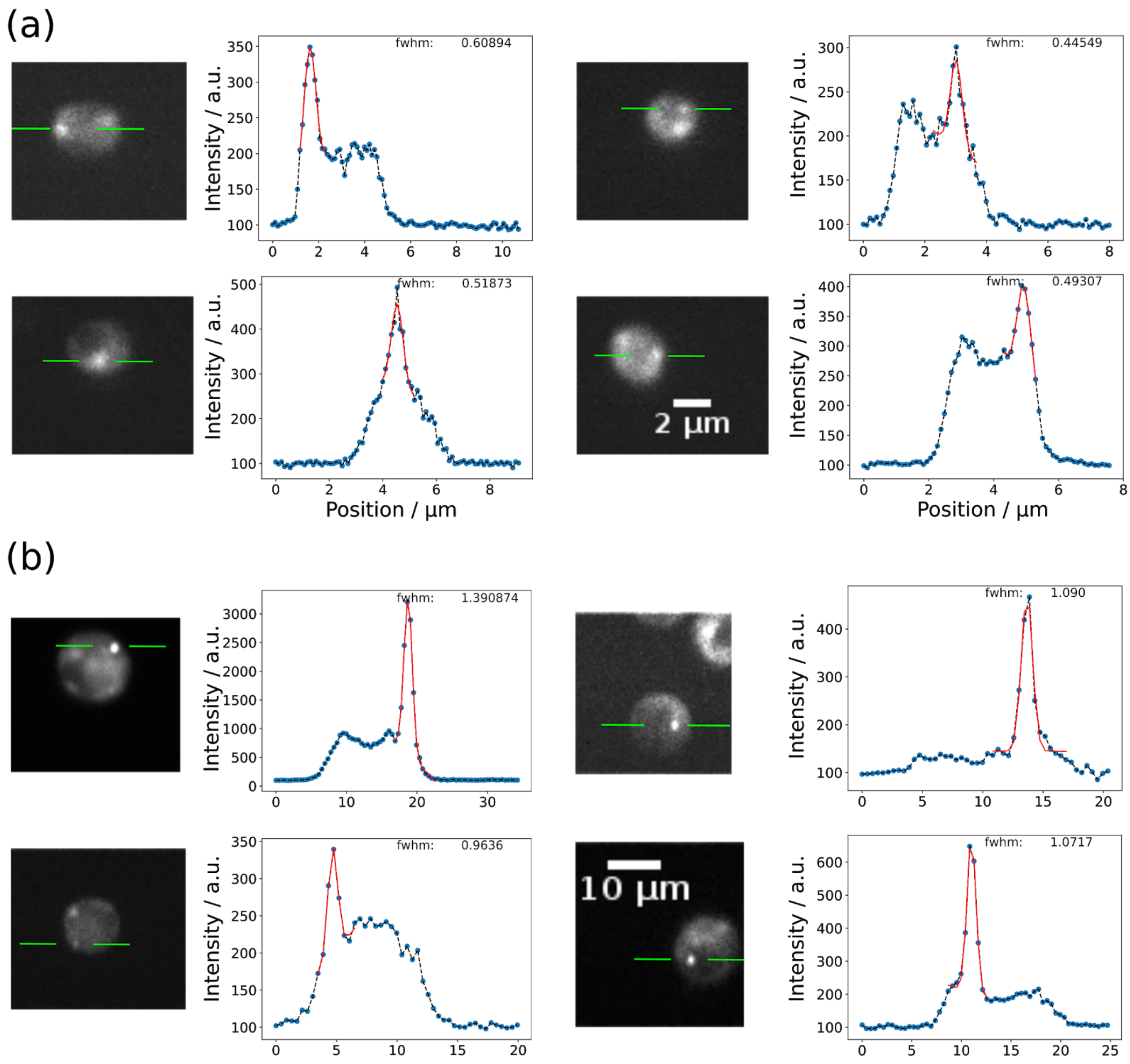


**Supplementary Fig. 6: Intensity line profiles of fluorescent P-bodies and stress granules.** The green lines in the images indicate the position of the intensity profile shown in the corresponding intensity plots (left part). The red line shows a Gaussian fit (mixed model consisting of a Gaussian and linear part to accurately model slanted background) to the intensity profile of a granule to extract full width at half maximum (FWHM). (a) Images of yeast cells recorded in flow using a 60X 1.2NA objective, with Cdc19-GFP stress granules exhibit a minimum FWHM around 500 nm. (b) Images of 293T cells recorded in flow using a 15X 0.5NA objective with mNG-AGO2 P-bodies exhibit a minimum FWHM around 1000 nm.

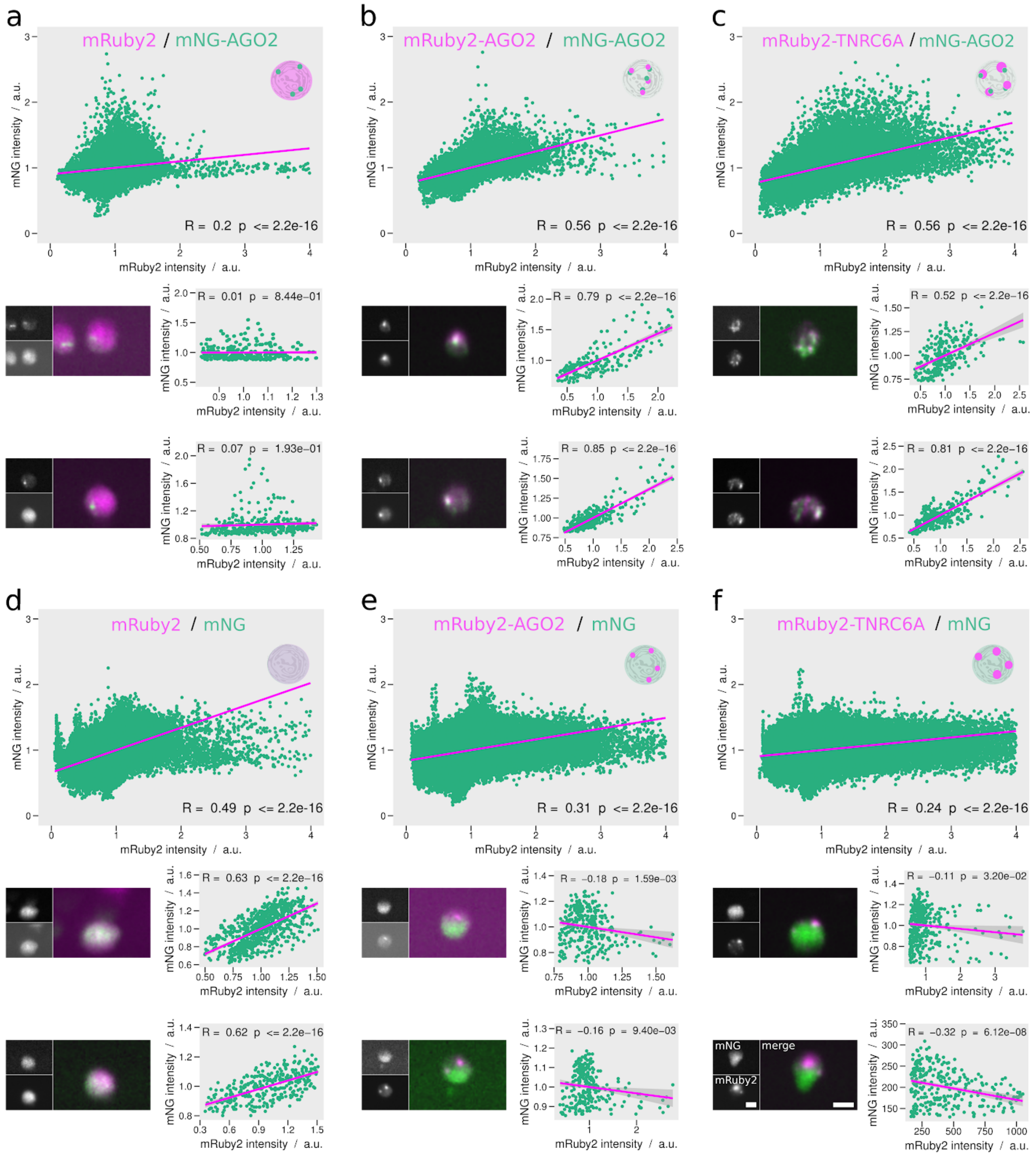


**Supplementary Fig. 7: Colocalization plots for single pixel intensities of normalized mNG and mRuby2 pixel intensities.** Main panels (a-f) display scatter plots of single pixel intensities from a population of approximately 1000 cells (top) as well as scatter plots of single pixel intensities and corresponding images of single cells (bottom) in the different co-expression situation as indicated. The Pearson correlation coefficient[3] is indicated for each correlation plot as a means to quantify the degree of colocalization between fluorophores. The pixel intensity for each cell was normalised to its average intensity value, with only the top 10% of the brightest pixels being retained for analysis. Low p-values in situations where co-localization between the two proteins is not expected (i.e. a, e, f) can be explained by the large number of data points (> 200000 points).[4] Scale bars are 10 µm.

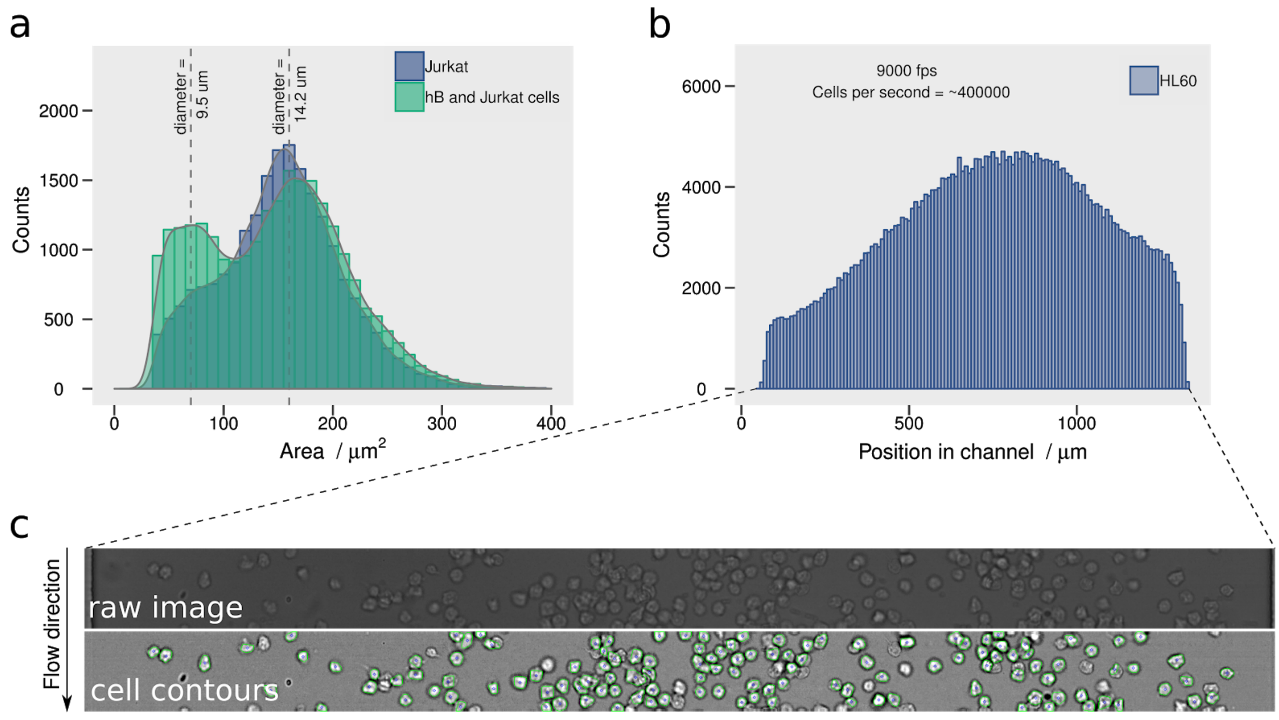


**Supplementary Fig. 8: Ultra-high-throughput brightfield imaging flow cytometry**. (a) Two different cell populations (a pure population of Jurkat cells and a mixture of Jurkat and Human B-lymphoid cells in a 1:1 ratio) were imaged in brightfield mode. The size histogram for the mixed cell population shows two distinct maxima, corresponding to the average diameters of the hB and Jurkat cells. (b) Cell distribution as a function of distance across the channel width ($w$ = 1331 µm, $h$ = 55 µm) at a camera frame rate of 9000 $s^{-1}$ and a shutter speed of 5 us, yields an experimental throughput of over 400,000 cells/second. (c) A single frame showing Jurkat cells in flowing at 0.25 m/s and the corresponding processed image, with the cell contours shown in green.

**Supplementary Note 1.**
Microfluidic device configuration.

The device schematic shown in Fig. 1b, consists of a single high aspect ratio (two loop) channel having a width of 665 µm and a depth of 59 µm. The use of only a single (wide) microfluidic channel avoids the need for evenly distributing sample across multiple channels. This removes imaging losses due to finite inter-channel spacing in the field of view and minimizes channel blockages due to small cross sections, leading to significant increases in analytical throughput. It is important to note that elasto-initial focusing provides control over the axial position of cells within the microfluidic channel and operates within a lower flow rate regime than related inertial focusing schemes. These features are ideal for high resolution imaging flow cytometry, since signal collection can be achieved using longer exposure times.

**Supplementary Note 2.**
Optical performance of the microfluidic imaging flow cytometer.

The sensor of the CMOS camera is read out line by line. Accordingly, any variation of one of the dimensions in the region of interest will significantly affect the maximum frame rate at which images can be acquired. To allow imaging of every single cell, we set the horizontal dimension to be slightly higher than the maximum diameter of a cell. Since we are only interested in a thin region of interest along the focused excitation line (the FWHM of the Gaussian laser beam is 15 µm), we were able to use this effect to increase the measurement frequency. Values for the ROI, based on the magnification of the objectives, are provided in the **Supplementary Table 1**. The effective exposure time is defined by the length of the excitation pulses, with longer pulses resulting in higher signals, but increased motion blur.

The resolving power of an optical system depends on the camera and associated pixel size. 6.5 µm pixels reduced in size by a 10x objective result in an effective pixel size of 0.65 µm, a 40X objective results in an effective pixel size of 0.16 µm and a 60X objective results in an effective pixel size of 0.108 µm. The 20X lens is useful for mammalian cells that adequate fit into the field of view. The pixel size using the 20X objective is 0.33 µm. The 40X objective with a pixel size of 0.16 microns provides a higher magnification for small objects and is the proper objective for imaging small sized cells such as yeast. The optimal operating speed in the imaging flow cytometer is approximately 0.05 m/sec that gives the best throughput. If combined with a 15X magnification this speed corresponds to the highest resolution setting (shown below) with a pixel size of 0.43 µm. Put simply a magnification of 20X in our system matches the pixel size of the Amnis system when a 60X objective is used.

According to established optical theory, optimal spatial resolution is achieved when matching the diffraction-limited resolution of the optical system to two pixels on the camera, in each linear dimension. This is commonly termed the Nyquist limit. Analysis of the specifications of the Amnis® ImageStream®X Mk II (Luminex, Austin, United States) indicates that objective magnifications cannot satisfy the Nyquist limit for achieving effective resolution, with the limit determining the maximum acceptable pixel size needed to meet the optical resolution of a specific objective. Given the need to fulfil the Nyquist criterion, the size of stress granules and P-bodies inside cells should be between 500 nm and 1 µm in diameter. For example, to resolve a 0.7 µm object, an objective (**Supplementary Table 1, 2)**, with a 20x magnification and an NA of 0.5 would be needed. At this magnification, the projected image would be 13.5 µm, and thus a camera with a pixel size of 6.7

µm (13.5/2) or smaller is necessary. This value matches closely with the pixel size of our CMOS camera.

**Supplementary Table 1.**
Performance characteristics of the commercial Amnis™ imaging flow cytometer reprinted from the official brochure.

| Magnification | 40X | 60X | 20X |
|---|---|---|---|
| Numerical Aperture | 0.75 | 0.9 | 0.5 |
| Pixel Size | 0.5 x 0.5 µm | 0.3 x 0.3 µm | 1 x 1 µm |
| Field of View | 60 x 128 µm | 40 x 170 µm | 120 x 256 µm |
| Imaging Rate | 2000 cells/s | 1200 cells/s | 4000 cells/s |

**Supplementary Table 2.**
Performance characteristics of the stroboscopic imaging flow cytometer.

| Magnification | 40X | 20X | 15X | 10X |
|---|---|---|---|---|
| Numerical Aperture | 0.75 | 0.5 | 0.5 | 0.5 |
| Pixel Size | 0.16 x 0.16 µm | 0.33 x 0.33 µm | 0.43 x 0.43 µm | 0.65 x 0.65 µm |
| Field of View | 332 x 25 µm | 665 x 25 µm | 887 x 25 µm | 1331 x 25 µm |
| Imaging Rate | 5350 cells/s | 10900 cells/s | 20500 cells/s | 61000 cells/s |

**Supplementary Note 3.**
Sensitivity of the microfluidic imaging flow cytometer.

We evaluated the sensitivity and detection range of our imaging flow cytometer using suspensions of fluorescent particles, characterized by four distinct fluorescence intensities. Based on the analysis of such calibration samples we are able to quantify our sensitivity, which is depicted in the calibration curve show in **Supplementary Fig. 4.** Absolute quantification of the fluorescence intensity in MESF (molecules of equivalent soluble fluorophore FITC) units was provided by the manufacturer (Quantum Alexa Fluor 488 MESF, Bangs Laboratories, Indiana, USA). Fluorescence intensities ranged between 3,179 and 333,766 MESF units. The smallest intensity peak in our calibration curve (population 1, green histogram: **Supplementary Fig. 4c**) has an intensity of 3,179 MESF units. This intensity lies below the calculated limit of detection (LOD) of the imaging cytometer, which corresponds to 8,481 MESF. Interestingly, we were still able to detect beads with an intensity of 3,179 MESF units by bandpass filtering the images and integrating pixels with an intensity above the background. It is also noteworthy that most applications in flow cytometry typically exhibit values above 30000 MESF[5].

**Supplemental Note S-4.**
Ultrahigh throughput of the microfluidic imaging flow cytometer.

After examining two different molecular weight PEO solutions over a wide range of flow rates, we concluded that focusing at the channel centerline remains efficient when using a low molecular weight PEO (1 MDa) solution at a flow rate between 0.01 and 0.1 m/s (**Supplementary Fig. 1d**). As previously discussed, the use of 500 and 1000 ppm 1 MDa PEO allows focusing of cells at flow velocities between 0.01 and 0.1 m/s, as shown in **Supplementary Fig. 1**. At a velocity of 0.1 m/s, cells only take 300 ms to cross the 30 µm wide field of view. This means that theoretically a minimum frame rate of 3,334 fps is required for accurate sampling. Using a 10x objective, a 1331 x 90 µm area can be imaged in the region of interest, yielding a throughput of 400,000 events per second, as shown in **Supplementary Fig. 8**. Accordingly, the imaging flow cytometer is able to image an enormous number of rapidly-moving cells in an ultra-high-throughput manner and with an imaging resolution defined by a 10x magnification objective.
